# Normative modeling of brain morphology reveals neuroanatomical heterogeneity and biological subtypes in major depressive disorder

**DOI:** 10.64898/2025.12.07.692810

**Authors:** Qingchen Fan, Jiahao Gao, Yihan Wu, Yun Wang, Ling Zhang, Jingjing Zhou, Yuan Feng, Yaxuan Lu, Gang Wang, Yuan Zhou

**Affiliations:** CAS Key Laboratory of Behavioral Science, Institute of Psychology, Beijing, China; Department of Psychology, University of Chinese Academy of Sciences, Beijing, China; State Key Laboratory of Cognitive Science and Mental Health, Institute of Psychology, Chinese Academy of Sciences, Beijing, China; Beijing Key Laboratory of Mental Disorders, National Clinical Research Center for Mental Disorders & National Center for Mental Disorders, Beijing Anding Hospital, Capital Medical University, Beijing, China; Advanced Innovation Center for Human Brain Protection, Capital Medical University, Beijing, China

**Keywords:** MDD, Normative modeling, Neuroanatomical heterogeneity, Biological subtypes

## Abstract

**BACKGROUND:** Major Depressive Disorder (MDD) is characterized by high neurobiological heterogeneity, which hinders precise diagnosis and treatment. Traditional group-level neuroimaging analyses fail to capture individual differences, while normative modeling offer a promising approach to quantify individual deviations from healthy brain structure patterns, facilitating the identification of biological subtypes and offering a data-driven framework to dissect this heterogeneity.

**METHODS:** Using 1,190 healthy controls, we constructed normative developmental trajectories of gray matter volume (GMV) across 246 Brainnetome-defined regions using Bayesian linear regression. Deviation maps were derived for 398 MDD patients. k-means clustering was employed to identify GMV-based biotypes. Then, the clinical characteristics and anatomical differences among these subtypes were explored, along with the post-treatment clinical features and treatment responses of participants who completed the 8-week antidepressant treatment within each subtype.

**RESULTS:** Patients with MDD exhibited widespread yet individually variable GMV deviations. Clustering analysis revealed two subtypes: Subtype 1 displayed predominantly negative deviations in sensorimotor and occipital cortices, whereas Subtype 2 showed widespread positive deviations in temporal and posterior cingulate regions. Subtype 1 had higher extraversion and symptom-linked deviation patterns; in Subtype 2, deviation burden correlated with generalized anxiety. Longitudinally, Subtype 1’s GMV deviation changes predicted symptom improvement, while Subtype 2’s deviations correlated with baseline severity.

**CONCLUSIONS:** Normative modeling of GMV reveals marked neuroanatomical heterogeneity in MDD and identifies subtypes with distinct clinical and treatment-related characteristics, laying a foundation for precision psychiatry and individualized interventions.

## INTRODUCTION

Major Depressive Disorder (MDD) has become a severely harmful mental illness worldwide, with its high prevalence and disability increasingly posing a major public health challenge[1,2]. MDD is primarily characterized by persistent low mood and anhedonia, but it exhibits substantial heterogeneity across clinical presentations, underlying neural mechanisms, and treatment responses. This poses significant challenges for disease diagnosis and treatment. Therefore, it is crucial to dissect heterogeneity in patients with MDD, which will provide a basis for formulating more targeted individualized treatment plans[3–6].

Neuroimaging techniques provide potential revenue to facilitate an understanding on heterogeneity in patients with MDD. Gray matter volume (GMV) derived from structural MRI, an easily-obtained and reproducible indicator reflecting brain structural characteristics, has been widely used to detect brain structural abnormalities in patients with MDD. However, traditional group-level analyses (comparing average GMV between MDD cohorts and healthy controls [HCs]) have yielded inconsistent findings: while some studies report reduced GMV in emotion-regulation regions (e.g., prefrontal cortex, cingulate gyrus, hippocampus)[7–9], others find no significant differences or even increased GMV in these same areas[10–12]. Even when consistent trends emerge at the group level (e.g., hippocampal volume reduction), the magnitude and regional distribution of GMV alterations vary drastically across individual patients[9,13]. The root of this inconsistency lies in the limitations of group-level paradigms: they aggregate data across heterogeneous patients, erase individual differences, and fail to treat “neuroanatomical heterogeneity” as a core variable to be investigated. As a result, these approaches cannot resolve the individual-specific structural anomalies that may underpin distinct MDD subtypes or pathological pathways[14].

To address these limitations and reduce the impact of heterogeneity, some studies have attempted to subtype MDD based on GMV and cortical thickness[15–18]. These subtypes may correspond to different biological mechanisms, facilitating the development of theories of “biomarker-based subtypes” or “neurobiological subtypes” and guiding personalized treatment. On the other hand, normative modeling, which has been proposed and widely applied in neuroimaging research on mental disorders, provides a new approach to advance understanding of heterogeneity in patients with MDD[13,14,19–22]. This approach involves constructing a “normal” distribution model of brain structure from large-sample healthy populations and using it as a benchmark to quantify individual deviations in brain imaging features for each patient. This method not only identifies which brain regions are abnormal but also quantifies the direction and degree of abnormalities, enabling quantitative characterization of individual brain structural developmental anomalies[21]. Compared with traditional methods, normative modeling significantly enhances the ability to resolve individual heterogeneity in neuroimaging data, particularly for highly heterogeneous diseases like MDD. This framework further enables linking individual-specific deviations to clinical symptoms, disease subtypes, and treatment responses[23]. Although several studies have applied this approach to explore the heterogeneity of MDD, existing research still faces certain limitations. Shao et al. constructed a normative model based on GMV to examine differential responses to antidepressant treatment across groups, but they did not employ individual deviation scores for subtyping[13]. Wu et al. built the normative model solely based on corpus callosum volume, which to some extent limited the understanding of widespread brain structural alterations in MDD[24]. Wang et al. constructed a normative model based on GMV and identified subtypes of MDD using individual GMV deviations; however, this work was limited to cross-sectional data and did not investigate longitudinal changes in gray matter volume deviations[15].

In this study, we employed sMRI data from 1,809 participants to comprehensively investigate the neurobiological heterogeneity and biological subtypes of MDD. We firstly established standardized growth models of GMV across 246 brain regions to characterize the neuroanatomical heterogeneity of MDD. Subsequently, through analyzing these deviations, we revealed interindividual heterogeneity in MDD patients and identified neurobiological subtypes based on their deviation patterns. Critically, a subset of patients underwent follow-up imaging after an 8-week course of antidepressant treatment, allowing us to examine longitudinal changes in GMV deviations and their relationship with symptom improvement. Moreover, we tested whether baseline deviation patterns could predict subsequent treatment outcomes. By integrating normative modeling, biotyping, and longitudinal follow-up, this study not only delineates the structural heterogeneity of MDD but also provides new insights into how brain deviation patterns are linked to therapeutic response, thereby advancing precision psychiatry for depression.

## MATERIALS and METHODS

### Participants

The datasets utilized in this study comprises two parts: a public dataset (Dataset I) and three hospital-based cohorts (Dataset II, Dataset III and Dataset VI) from Beijing Anding Hospital. For Dataset I, we used a publicly available dataset Human Connectome Project (HCP) S1200 release dataset, which has 1,113 subjects (507 males; mean age, 28.8±3.6; age range, 22-37) (http://www.humanconnectome.org/). All the scans and data from the individuals included in the study had passed the HCP quality control and assurance standards. Dataset I was used to provide a large healthy reference sample for constructing the normative modeling. For Dataset II-VI, the three hospital-based cohorts comprised 290 healthy controls and 406 patients with MDD. The healthy controls from these hospital-based cohorts were combined with Dataset I to construct the normative model (Fig S1b), while the patients with MDD were used to investigate GMV deviations relative to the normative reference. For Dataset II, 67 patients with depression received 8-week antidepressant treatment prescribed by professional psychiatrists according to their symptoms after recruitment. Among them, 46 patients were treated with SSRIs (escitalopram, sertraline, fluoxetine, or fluvoxamine), 2 patients received SNRIs (venlafaxine or duloxetine), and the remaining 19 patients were treated with other antidepressants (such as bupropion and Biqi Tian oligosaccharide capsules). Among them, 37 patients received SSRIs treatment and underwent imaging data collection at the end of week-8. Details of inclusion and clinical assessment of patients are provided in Supplementary Method 1. The current study was approved by the Human Research and Ethics Committee of Beijing Anding Hospital, Capital Medical University, and all participants provided signed informed consent.

### MRI Data Acquisition and Processing

All structure MRI in the hospital-based cohorts were acquired using a Siemens 3T-MRI scanners. The detailed scanning parameters are listed in Table S1. We utilized FreeSurfer data from the 1,113 healthy controls in the HCP dataset (Dataset I), which included structural volumes. These data were preprocessed using the FreeSurfer pipeline (version 5.3.0) and are publicly available through the HCP. For Datasets II–IV, we applied the same FreeSurfer pipeline (version 5.3.0) to derive structural volumes from T1-weighted images. The analysis was conducted using the “recon-all” command[25]. After preprocessing the T1-weighted images, the GMV for each participant were quantified via the Brainnetome Atlas[26]. This approach enabled us to calculate GMV for 210 cortical subregions and 36 subcortical structures, resulting in a total of 246 distinct neuroanatomical structures (Figure 1A). Finally, we conducted data screening to exclude participants with poor-quality data and those who were either too young or too old, ensuring that the age-related trajectories of GMV constructed from the HC could adequately cover the age range of the MDD[27,28] (Table 1). For the HC group, participants with GMV outliers were excluded[27](Supplementary Method 2). Ultimately, 398 depressed patients and 1190 HCs were enrolled in this study (Fig S1c).

**Figure 1.**
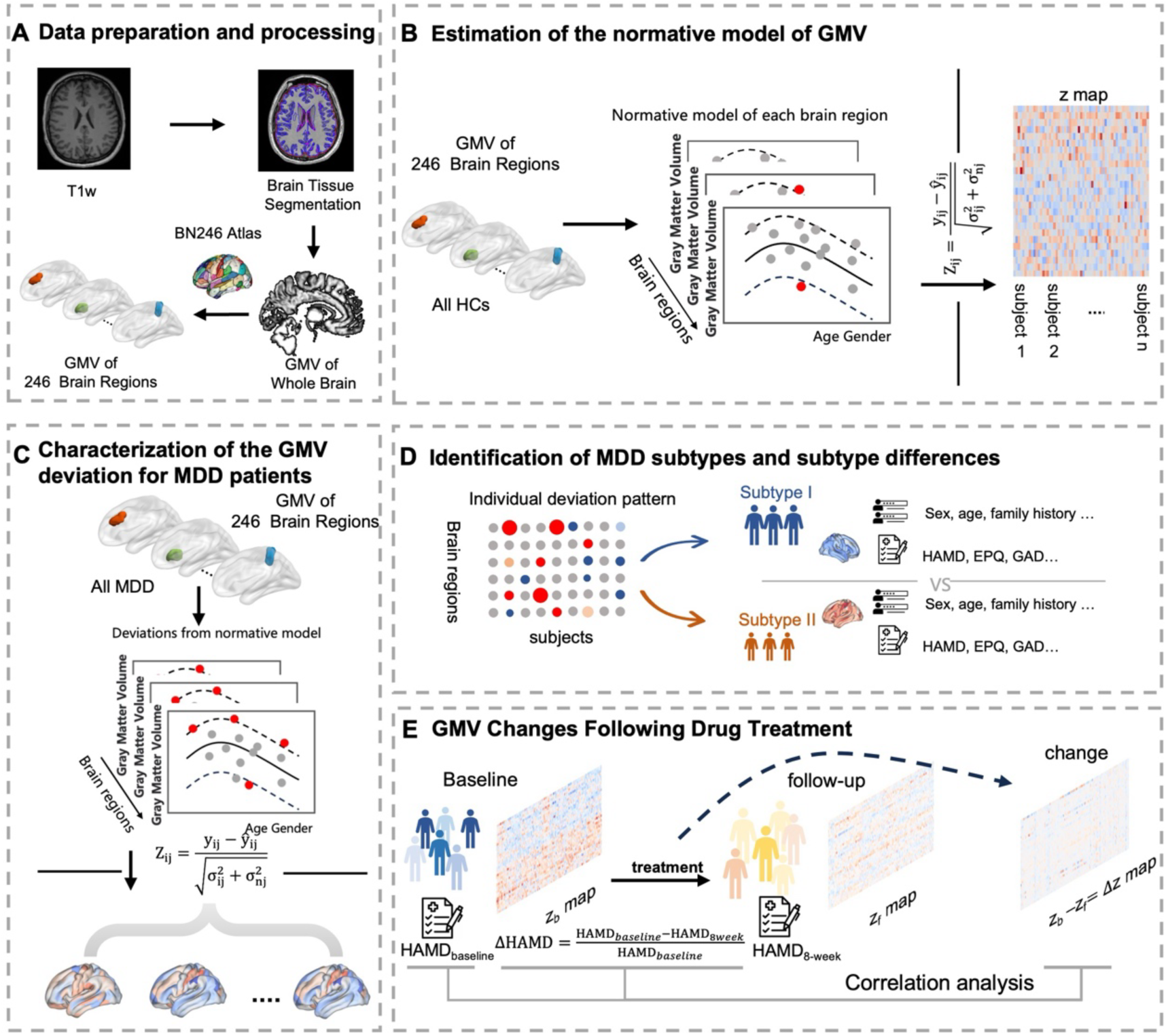
Overview of Study Workflow. **A** Brainnetome atlas comprising 246 brain regions (210 cortical subregions and 36 subcortical nuclei) used for GMV extraction. **B** Schematic of the normative model construction. Bayesian linear regression was used to establish age-related GMV trajectories in HCs. Individual GMV values are then projected onto these trajectories to calculate z-score deviation maps. **C** Generation of individual deviation (Z-score) maps for each patient with MDD by comparing observed GMV against the normative model. **D** Data-driven subtype identification using k-means clustering on deviation maps, followed by clinical and longitudinal analyses. **E** For the longitudinal analyses, deviation maps were computed at baseline and after 8 weeks of antidepressant treatment. The Δ z maps representing within-subject changes were modeled using Support Vector Regression and Partial Least Squares to assess their associations with clinical improvement (ΔHAMD) and baseline depressive severity.

**Table 1.** Clinical characteristics of the participants.

| DataSet | Group | Age range | Age, Years,<br>Mean (SD) | Sex, Male /Female,<br>n | First Episode/Recurrent<br>Episode, n | Family History of<br>Psychiatric Disease,<br>Yes/No | HAMD-17,<br>Mean(SD) | SSRIs/SNRIS/<br>Others/No |
| --- | --- | --- | --- | --- | --- | --- | --- | --- |
| Data I | HCs, n = 993 | 22-37 | 28.97 ± 3.69 | 395/598 | - | - | - | - |
| DataII | Patients, n = 174 | 19-37 | 26.41 ± 4.99 | 53/121 | 82/92 | 34/140 | 20.62 ± 4.17 | 127/8/31/14 |
|  | HCs, n = 89 | 19-37 | 24.66 ± 4.16 | 29/60 | - | - | - |  |
| | t or $\chi^2$, p | - | t = 4.00,<br>p = 0.000 | $\chi^2$ = 0.044<br>p = 0.832 | - | - | - | |
| DataIII | Patients, n=114 | 19-37 | 26.40 ± 4.49 | 36/79 | 66/45(-4)*<sup>1</sup> | - | 20.96 ± 4.35 | 0/0/0/115 |
|  | HCs, n= 66 | 20-37 | 26.68 ± 4.20 | 10/56 | - | - |  |  |
| | t or $\chi^2$, p | - | t = -0.40,<br>p = 0.686 | $\chi^2$ = 4.951<br>p = 0.026 | - | - | | |
| DataIV | Patients, n=110 | 19-37 | 27.10 ± 4.48 | 31/80 | 43/66(-2)*<sup>2</sup> | 13/98 | 20.79 ± 3.66 | 12/0/3/96 |
|  | HCs, n= 42 | 20-36 | 26.05 ± 3.97 | 12/30 | - |  |  |  |
| | t or $\chi^2$, p | - | t = 1.34,<br>p = 0.183 | $\chi^2$ = 0.044<br>p = 0.832 | - | | | |

\*1 In DataIII, 4 cases lack statistical records on whether the subjects are first-episode or recurrent.
\*2 In DataIV, 2 cases lack statistical records on whether the subjects are first-episode or recurrent.
SSRIs: Selective Serotonin Reuptake Inhibitors
SNRIs: Serotonin-Norepinephrine Reuptake Inhibitors

### Constructing a Normative Modeling of GMV

Using the PCNtoolkit (https://pcntoolkit.readthedocs.io/)[29]with Python 3.6 and data from healthy participants in Datasets I and II–IV, we randomly divided the sample into training (n×80% = 991) and testing (n×20% = 199) sets. Based on the training set, normative developmental trajectories of GMV (gray matter volume) were established for each brain region as a function of age (Figure 1B). A Bayesian Linear Regression (BLR) model[30] was employed to characterize regional GMV as a nonlinear function of age, while adjusting for confounding factors including scanner site effects, sex, and total intracranial volume (TIV). This framework incorporated multicenter data heterogeneity through hierarchical priors and allowed flexible modeling of both linear and nonlinear relationships[31–33]. The model’s generalizability was evaluated via 10-fold cross-validation, with the model fit for each region.

Subsequently, regional z scores were computed in the testing set of healthy participants to quantify individual deviations from the normative range. The z score was defined as the difference between the observed GMV and the model-predicted value, normalized by model uncertainty (Formula 1). After model validation, the same procedure was applied to patient data to assess deviations of GMV from normative developmental trajectories.

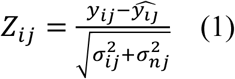

where is the true GMV, is the predicted mean, is the estimated noise variance (uncertainty in the data), and is the variance from modeling uncertainty for each participant (i) and each brain region (j).

These z scores were integrated into individual deviation maps, providing a spatial visualization of neuroanatomical deviations from population-level age-related trajectories of gray matter changes. The individual deviation maps framework provides individualized quantification of localized GMV abnormalities, serving as a data-driven approach for identifying patient-specific neurostructural alterations.

### Estimation of individual GMV deviation patterns in normative modeling

To assess deviations in GMV among individuals with MDD relative to healthy individuals, we first mapped the GMV of each brain region in MDD patients from Dataset II-VI to a normative percentile atlas derived from HCs (Figure 1C). This allowed us to quantify the deviation of each participant’s GMV from the normative modeling. Then we calculated the z scores representing the deviation of GMV for each brain region in MDD patients. To define extreme individual-level deviations in GMV, brain regions with |Z|>1.96 (i.e., exceeding the two-tailed 95% CI) were classified as exhibiting extreme deviations (positive or negative). Subsequently, we performed two-sample t-tests to compare between-group differences in the total number of brain regions with extreme deviations (Total Outlier Count, TOC)[34], as well as the degree of deviation (Modulus)[23] between the MDD groups and the healthy controls in the testing set. Significance was corrected for multiple comparisons using the false discovery rate (FDR) method, with a corrected threshold of q < 0.05 (Supplementary Method 3). To evaluate inter-individual heterogeneity within the MDD group, we computed the overall deviation degree for each participant and analyzed the spatial overlap patterns of extreme deviations across individuals.

### Identifying MDD Biotypes Based on Individual GMV Deviations

To identify MDD subtypes with distinct deviation patterns, we employed a data-driven k-means clustering algorithm (Figure 1D). In cluster analysis, individual deviation maps were used as features, and a similarity matrix was created by calculating the Euclidean distance between them. For each predefined cluster number *k* (*k* ∈ [2,10]), the clustering was repeated 10 times with different random seeds to reduce sensitivity to initial conditions. The optimal k was determined using an ensemble strategy: the NbClust package[35] calculated 30 clustering validity indices, and the k supported by the majority of indices was selected as the final result.

### Clinical Characteristics of MDD Biotypes

For the identified subtypes, we firstly compared their differences in GMV deviations and clinical variables. Subsequently, we applied multivariate Partial Least Squares (PLS) analysis for each biotype to investigate the covariance structure between brain deviations (X-variables) and clinical symptoms (Y-variables). The statistical significance of the extracted latent variables was assessed using permutation tests (n = 1,000) to determine whether these components explained covariance exceeding random chance. For significant PLS components, Pearson correlations between X-scores (latent scores of brain deviations) and Y-scores (latent scores of symptoms) were calculated to represent the shared variance between neuroanatomical features and clinical measures. The significance of these correlations was further confirmed through additional permutation tests (n = 1,000) (Supplementary Method 4).

### Association between Individual Deviation Map Changes and Depressive Symptom Scores Before and After Treatment

For patients in both subtypes who completed 8 weeks of treatment (N_subtype1_ = 20, N_subtype2_ = 17), we first computed the individual deviation maps of GMV at week-8 by mapping each patient’s regional GMV values to the normative percentile atlas derived from HCs. Second, we calculated the individual deviation maps change by subtracting the week-8 deviation map from the patient’s baseline deviation map. Then the individual deviation map changes were mapped to the 7 networks and subcortical regions. Subsequently, support vector regression was employed to explore the correlation between the network-level changes in individual deviation maps and the concurrent changes in HAMD scores within each subtype. To ensure model robustness, we applied repeated two-fold cross-validation, with prediction accuracy indexed by the Pearson correlation between predicted and observed HAMD changes. Statistical significance was assessed using 1,000 permutations, and model weights were further analyzed to identify key brain networks contributing to prediction.

Additionally, to investigate the potential impact of baseline depressive symptom severity on treatment-related changes in GMV within each subtype, we employed PLS to examine the relationship between changes in individual deviation maps and baseline HAMD scores (Figure 1D). The statistical significance of PLS components was evaluated using permutation tests (n = 1,000). For each significant component, we extracted individual component scores and computed their Pearson correlation with baseline HAMD scores, with significance again evaluated using permutation tests (n = 1,000) (Supplementary Method 5).

## RESULTS

### Normative growth models of GMV reveal remarkable neuroanatomical heterogeneity in MDD

We constructed normative modeling for GMV in 246 brain regions. Between-group analysis revealed significantly greater GMV deviations in the MDD group, including both the number of deviant brain regions (TOC) (Cohen’s d = 0.49, FDR-corrected q <0.01) and the degree of deviation (Cohen’s d = 0.54, FDR-corrected q <0.01) (Figure 2A). Regional analysis revealed positive deviations in the ventromedial prefrontal cortex, angular gyrus, and right middle temporal lobe and negative deviations in occipital lobe and sensorimotor regions in MDD patients compared to HCs (Figure 2B). Notably, at the individual level, 88.4% (n = 351) of patients demonstrated extreme positive deviations in at least one brain region, while 80.4% (n = 320) showed extreme negative deviations in at least one region (Figure 2C). However, the prevalence of abnormal deviations in specific individual brain regions displayed marked heterogeneity: for any single brain region, no more than 10.1% (n = 40) of patients exhibited extreme positive deviations, and no more than 9.54% (n = 38) exhibited extreme negative deviations (Figure 2D). Collectively, while the vast majority of patients exhibited extreme GMV alterations in some brain regions, the specific affected regions demonstrated substantial interindividual variability.

**Figure 2.**
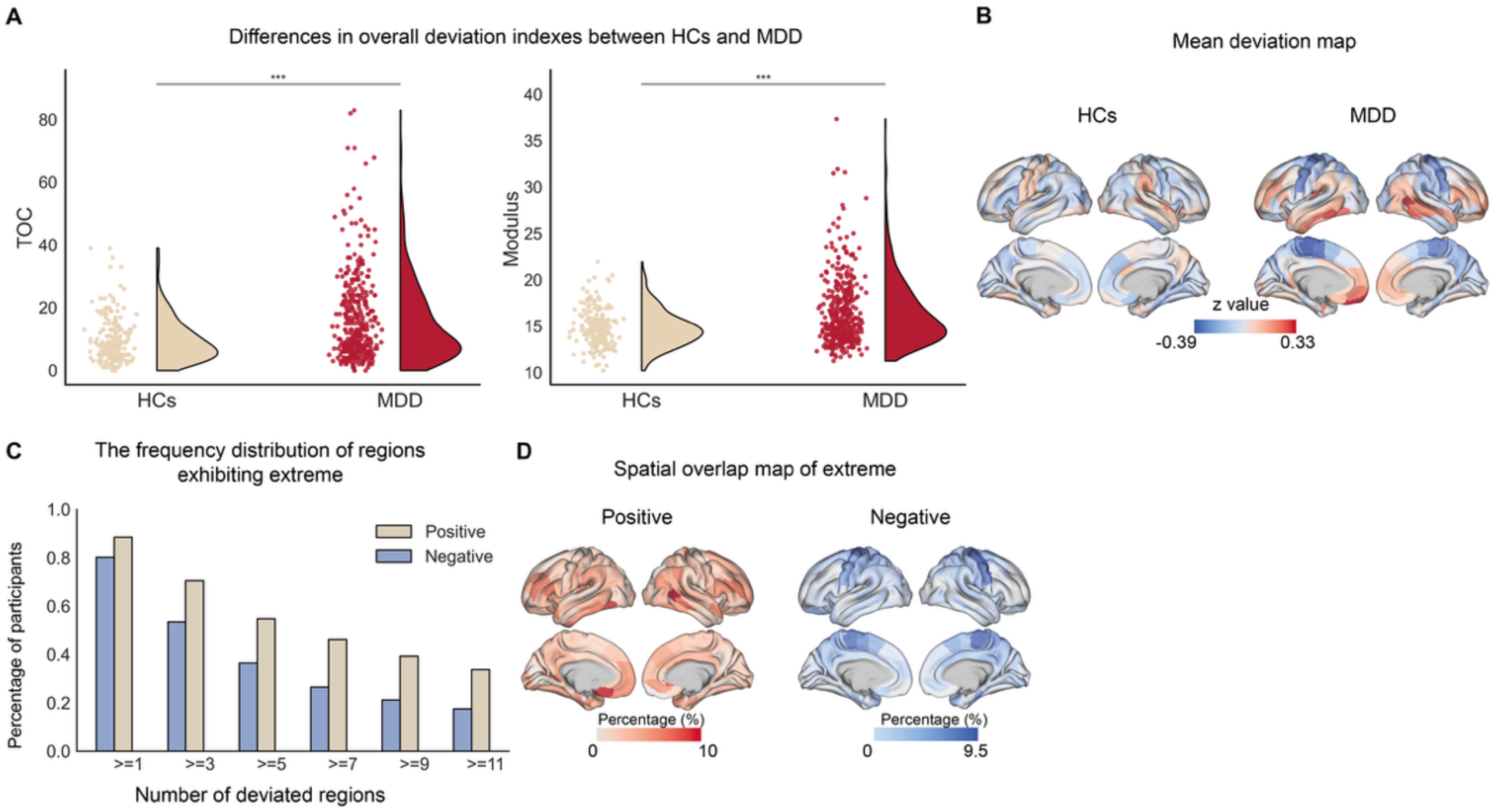
Neuroanatomical heterogeneity in major depressive disorder revealed by normative modeling of gray matter volume. **A** Patients with MDD (n = 398) show significantly higher TOC and Modulus compared to healthy controls (n = 1,190; Cohen’s d = 0.49 and 0.54, respectively; FDR-corrected q < 0.01). **B** Spatial patterns of mean GMV deviations (blue, negative deviations; red, positive deviations). C Bar plots show the distribution of the number of regions per patient with extremely positive (pale yellow) and negative (blue) deviations. **D** Spatial overlap maps of extreme deviations. The color bar indicates the percentage of participants exhibiting extreme positive (left) or negative (right) deviations in specific regions.

### Data-Driven Identification of Two MDD Subtypes with Divergent Brain GMV Deviation Patterns

Using the k-means method, two MDD subtypes (Subtype 1 = 236, Subtype 2= 162) with different deviation patterns were identified. The optimal number of subclusters was consistently supported by 15 out of 23 effective quality indices (Figure 3A). We used the t-SNE dimensionality reduction algorithm to visualize the distribution of the two subtypes in a low-dimensional feature space. The results show that Subtype 1 and Subtype 2 are clearly separated in the two-dimensional space, indicating significant differences between the two subtypes in the feature space (Figure 3B). For the two subtypes, mapping the deviation values to the heat map allows for the intuitive observation that the overall pattern exhibits opposite deviation patterns (Figure 3D). Subtype 1 exhibited a predominantly negative deviation pattern across most brain regions, specifically in sensorimotor, occipital lobe and posterior cingulate gyrus, with minor positive deviation in prefrontal lobe. Subtype 2 showed the opposite deviation pattern, characterized by predominantly positive deviations in most brain regions, such as temporal lobe, posterior cingulate gyrus, and minor negative deviations in central sulcus (Figure 3C). Statistical comparisons between subtypes revealed significant differences in two indices: the number of deviant brain regions and the degree of deviation. We found that the overall number of deviant brain regions in Subtype 1 was lower than that in Subtype 2 (Cohen’s d = −0.95, FDR-corrected q < 0.01). Similarly, the degree of deviation in Subtype 1 was also lower than that in Subtype 2 (Cohen’s d = −0.90, FDR-corrected q < 0.01) (Figure 3E F).

**Figure 3.**
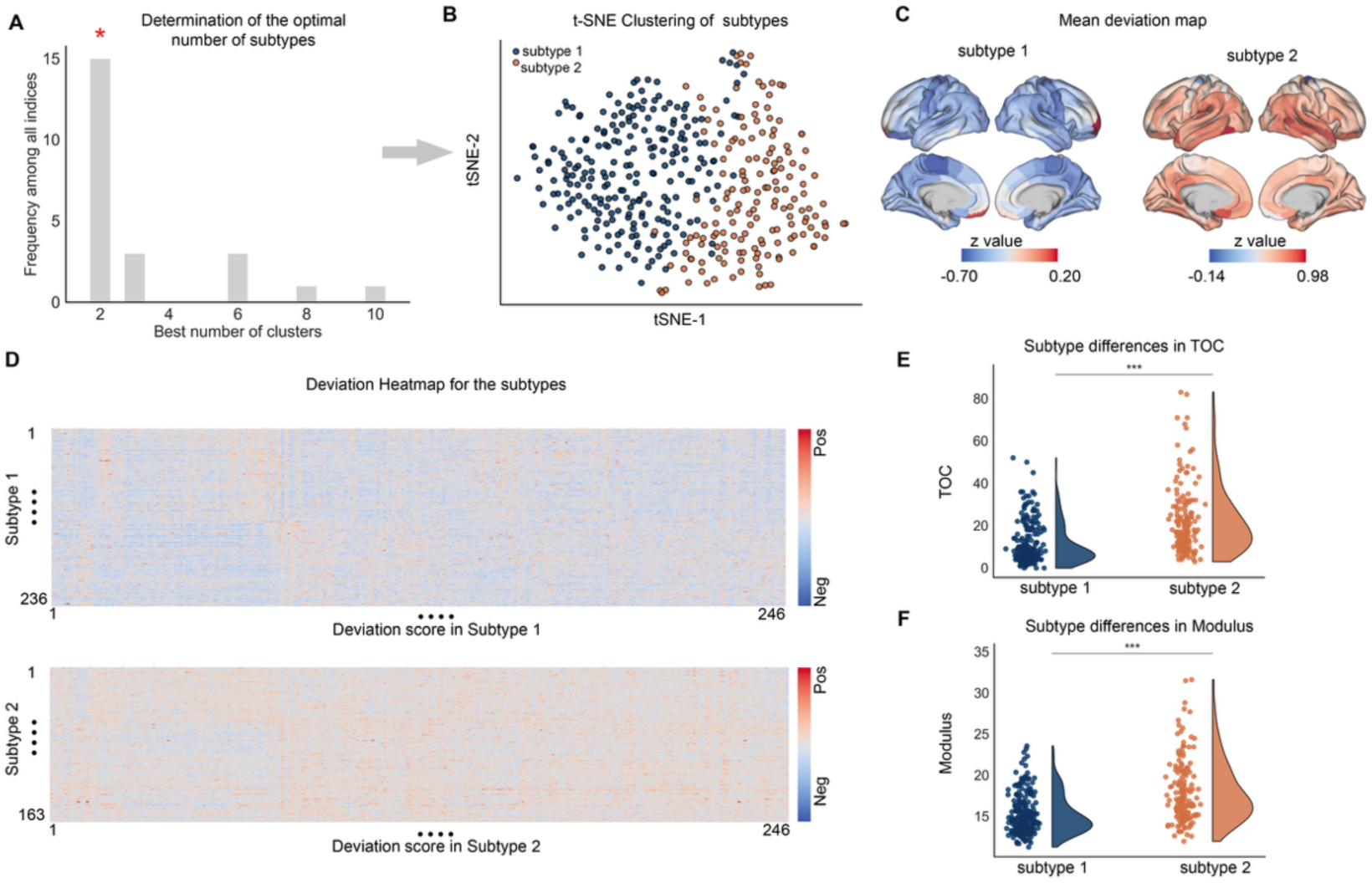
Two MDD subtypes with distinct neuroanatomical deviation patterns. **A** Determination of the optimal number of clusters. **B** t-SNE visualization of the two identified subtypes, showing clear separation between Subtype 1 (blue) and Subtype 2 (orange) in the feature space. **C** Spatial patterns of mean GMV deviations for the two subtypes (blue, negative deviations; red, positive deviations). **D** Heatmaps show opposite GMV deviation patterns. **E** Comparison of the TOC between subtypes. **F** Comparison of the deviation Modulus.

### Two Neurobiological Subtypes of MDD Differences in Clinical Characteristics

Significant differences were observed in sex distribution (Subtype1 M:F = 55:181; Subtype2 M:F=64:98; *x*^2^ = 11.26 *p* = 0.001). And Subtype1 demonstrated higher extraversion scores than Subtype 2(Cohen’s d=0.43, p < 0.05). No other differences were detected in age, HAMD scores, or scores on other personality dimensions (Figure 4A, Table S2). Within each subtype, the clinical correlates of the TOC were explored. Only the total number of extreme deviation brain regions was positively correlated with GAD scores in Subtype 2 (r = 0.36, p < 0.05), with no such correlation in Subtype 1 (r = −0.05, p = 0.32) (Figure 4B).

**Figure 4.**
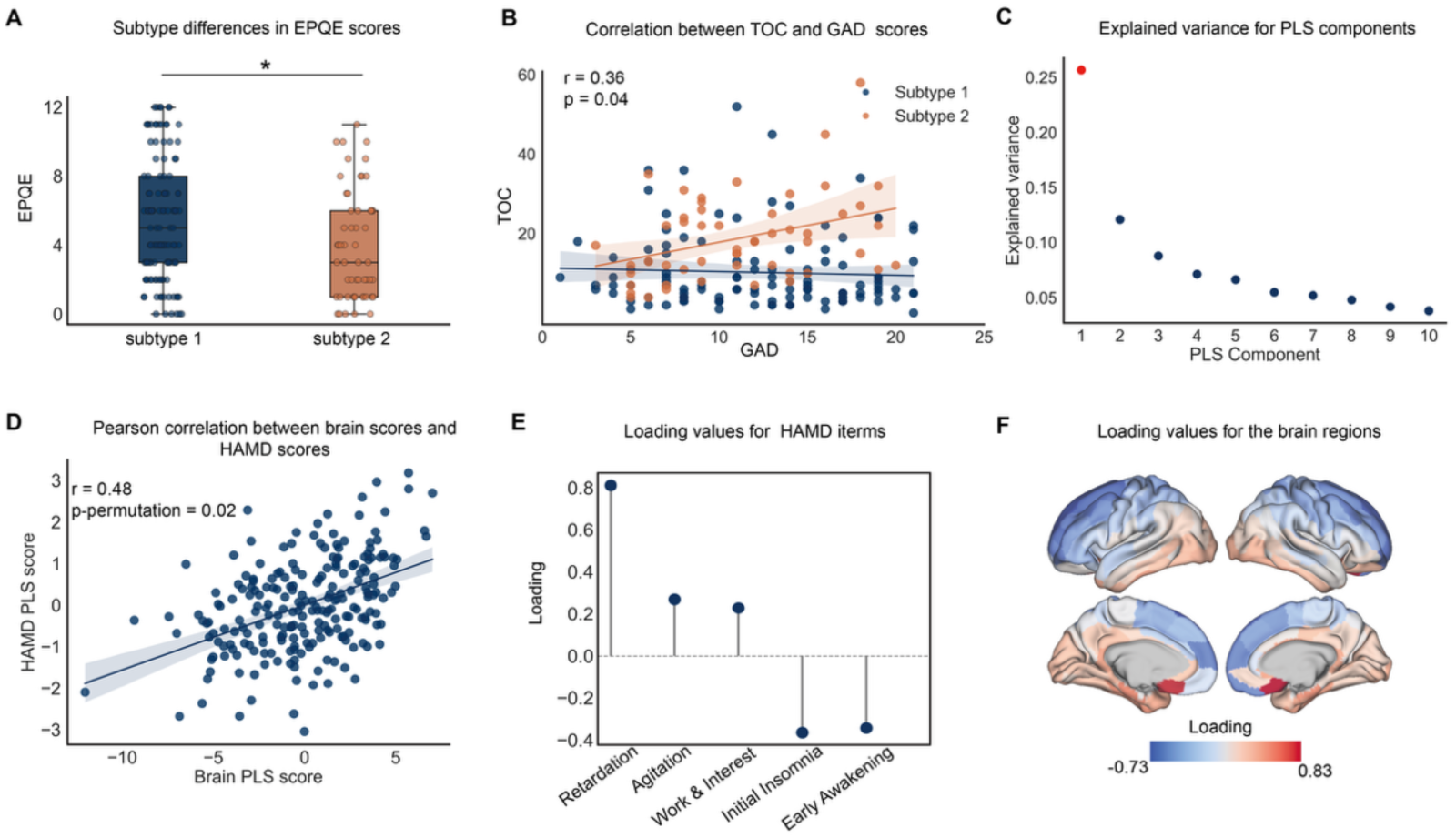
Clinical characteristics and symptom associations of the MDD subtypes. **A** Differences in personality traits between subtypes. Subtype 1 exhibited significantly higher Extraversion scores (EPQE) compared to Subtype 2. Cohen’s d=0.43, *p < 0.05 **B** Scatter plot showing the correlation between TOC and GAD scores. **C** Explained variance of PLS components in subtype 1. The significant PLS component is represented in red. **D** Pearson correlation between brain deviation scores and HAMD symptom scores in Subtype 1. **E** Loading values for specific HAMD items in Subtype 1.**F** Brain region loadings for the significant PLS component in Subtype 1.

In Subtype 1, the first component of the PLS model explained 26% of the variance in HAMD scores (permutation test p-value < 0.01; Figure 4C), with brain region scores showing a significant positive correlation with symptom scores (r = 0.48, permutation test p = 0.02; Figure 4D). The contribution loadings showed the highest positive values for retardation, agitation, and work-interest and negative values for initial insomnia and early awakening (Figure 4E). Positive loadings of brain structural deviations were primarily located in lateral and medial temporal cortices and cingulate cortices, while negative loadings were found in the sensorimotor cortices and superior frontal gyrus (Figure 4F). However, no association between brain structural deviations and HAMD scores was observed in Subtype 2.

### GMV Changes Following Drug Treatment

For Subtype 1 participants, the change in individual deviation maps from baseline to week-8 robustly predicted the corresponding change in HAMD-17 scores (permutation test p < 0.05) (Figure 5A). The features contributing most positively were localized to the subcortical network and visual network, whereas the strongest negative contributions were observed in the frontoparietal network and dorsal attention network (Figure 5B). In contrast, individual deviation maps change from baseline to week-8 did not directly predict HAMD-17 score changes in Subtype 2. However, multivariate analyses examining the relationship between these deviation-map changes and baseline HAMD-17 scores in subtype 2 revealed that the first latent component of a PLS model explained 38% of the variance in baseline HAMD-17 scores (permutation test p < 0.01; Figure 5C). Brain scores derived from this component were strongly and positively correlated with symptom scores (r = 0.91, permutation test p = 0.03; Figure 5D). Variables related to work-interest, depression, and weight contributed positively to this association, whereas early awakening and initial insomnia exhibited negative loadings (Figure 5E). Positive structural deviation changes were localized primarily to the angular gyrus, temporal lobe, and right ventromedial prefrontal cortex, whereas negative loadings were found in the sensorimotor cortex and posterior cingulate cortex (Figure 5F). Notably, these multivariate associations were absent in Subtype 1.

**Figure 5.**
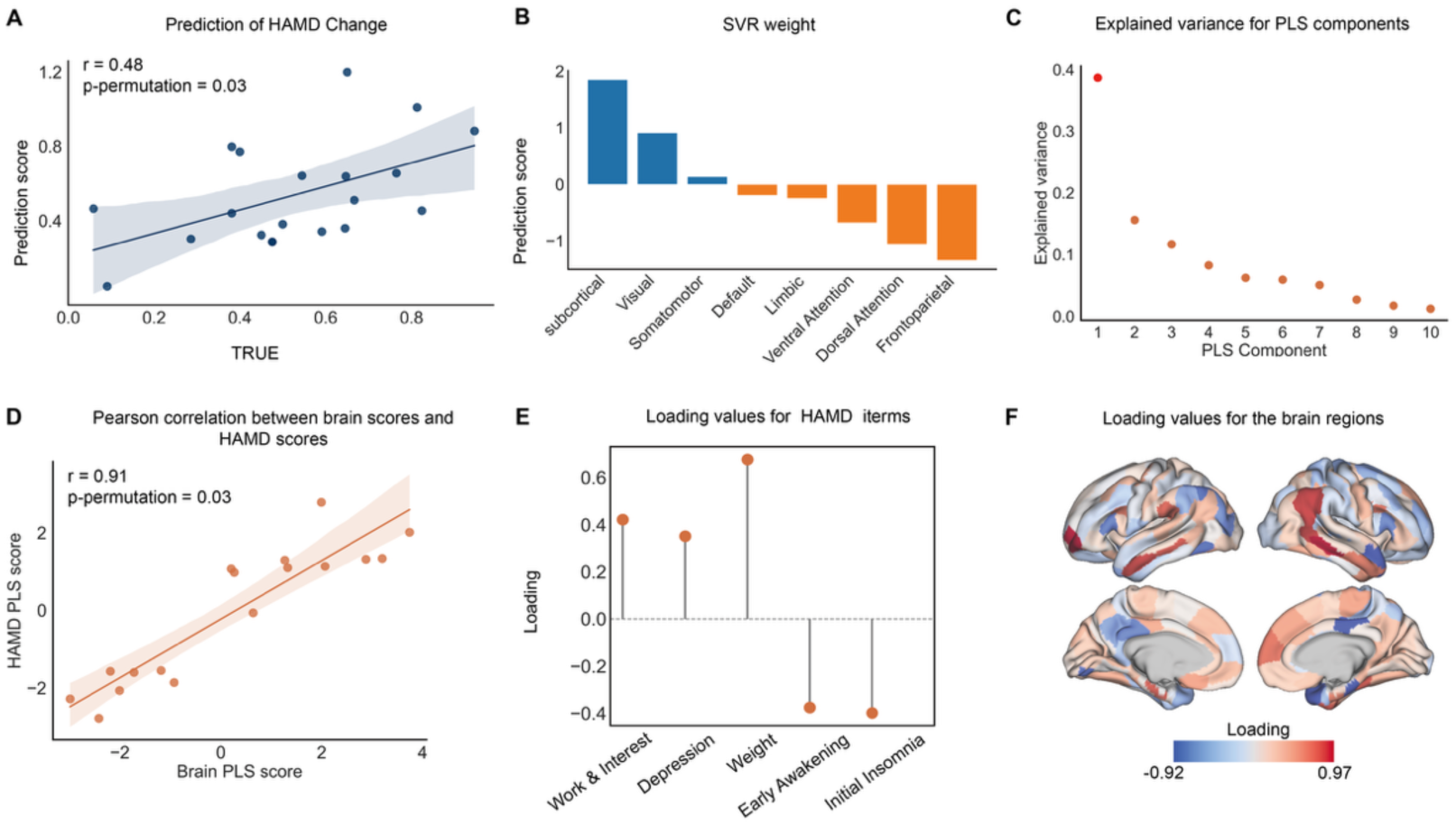
Longitudinal GMV deviation changes and their relationship to treatment outcomes. **A** The change in individual deviation maps from baseline to week-8 significantly predicted the change in HAMD scores in subtype 1. **B** SVR weights for Subtype 1, indicating that positive contributions from subcortical and visual networks and negative contributions from frontoparietal and dorsal attention networks drove the prediction of symptom improvement. **C** Explained variance of PLS components in subtype 2. The significant PLS component is represented in red. **D** Pearson correlation between brain deviation changes scores and baseline HAMD symptom scores in Subtype 2. **E** Loading values for HAMD items in Subtype 2. **F** Brain region loadings for the significant PLS component in Subtype 2.

## DISCUSSION

Normative modeling of GMV in this study quantified individual deviations from healthy brain structure patterns, revealing widespread yet highly variable neuroanatomical anomalies in MDD. Using these individual deviation profiles as features, unsupervised clustering identified distinct subtypes with divergent GMV deviation patterns. These subtypes differ in clinical characteristics, highlighting the biological heterogeneity of MDD. This approach demonstrates the capability of normative modeling to capture individual variability, offering a data-driven framework for dissecting the complexity of MDD and facilitating neurobiologically informed subtyping and mechanistic research.

### Normative Modeling Reveals Neuroanatomical Heterogeneity in Depression

Gray matter abnormalities are indeed a core neuropathological feature of depression[36]. Normative modeling utilizes large samples of healthy individuals to establish "growth curves" of brain structural development, enabling statistical inference of deviations between an individual’s GMV and the expected normative range. Compared to traditional group comparisons, this approach is well-suited to capturing highly individualized abnormal patterns in psychiatric disorders[37–39]. For instance, Shao et al. [13]demonstrated that, when applying a GMV-based normative model, structural abnormalities in patients with MDD were highly individualized—fewer than 11% of patients exhibited extreme deviations in any single brain region—suggesting an absence of universal regions of abnormality. Our findings similarly show that most patients displayed both positive and negative deviations across different regions, with no single brain area showing consistent abnormalities across the majority of patients. These results further underscore the anatomical heterogeneity of MDD. However, despite its strength in extracting rich individual-level information, normative modeling often yields high-dimensional outputs that are difficult to interpret directly[40]. Thus, multivariate techniques such as dimensionality reduction or clustering are necessary to map these deviations onto latent factors or subtypes, thereby facilitating the biological interpretation of complex individual differences.

### Data-Driven Subtyping of MDD and Its Clinical Relevance

In this study, we applied k-means clustering to individual GMV deviation maps and identified two distinct neuroanatomical subtypes of MDD. Subtype 1 was characterized by a pattern of negative deviation in GMV across brain regions, whereas Subtype 2 exhibited a pattern of positive deviation. This mirror-like pattern of structural deviation aligns closely with previous findings. For example, Chen et al. [18]identified two GMV-based subtypes in MDD: one showing gray matter increases in the frontal, parietal, and temporal lobes, and the other showing decreases in the thalamus, cerebral lobes, and limbic system. Similarly, Liu et al. [6]using multilayer functional connectivity and normative modeling, discovered two functional subtypes—one with positive deviations in the prefrontal/default mode network and negative deviations in the occipital/sensorimotor network, and the other showing the opposite pattern. These studies consistently suggest that both structural and functional brain measures can uncover biologically distinct subtypes within the heterogeneous depression population. In our analysis, each subtype exhibited relatively homogeneous deviation patterns within the group and marked divergence between subtypes, implying potential differences in underlying neurobiological mechanisms.

More importantly, such data-driven subtypes often show distinct clinical profiles and treatment responses[13,41,42]. For instance, Wang et al. [43] used clustering of functional connectivity and symptoms to delineate an insomnia-dominant subtype and an anhedonia-dominant subtype, which differed significantly in brain network profiles and clinical features. Likewise, Drysdale et al. [41] identified four “neurobiological subtypes” of depression based on resting-state connectivity and showed that these subtypes predicted differential responses to transcranial magnetic stimulation. In our study, the two subtypes differed in personality traits (e.g., extraversion). Furthermore, multi-omics studies have shown that imaging-based subtypes may also correspond to distinct genetic and immune profiles. For example, Tang et al.[44] reported that three imaging-defined subtypes were enriched in pathways related to neurodevelopment, inflammation, or lacked identifiable biomarkers. Taken together, identifying neuroimaging subtypes of depression offers a promising avenue for understanding its biological heterogeneity and may inform precision diagnosis and targeted interventions.

### Longitudinal GMV Deviations and Treatment Response

Our results show that changes in GMV deviation patterns predicted concurrent changes in HAMD scores only in Subtype 1, with the strongest positive contributions observed in subcortical and visual networks. These findings, consistent with previous research[45,46], suggest that alterations in brain areas implicated in emotion regulation and sensory processing are key to treatment response. Notably, the subcortical regions, which are involved in emotional processing, have long been associated with MDD, where alterations in areas like the amygdala and hippocampus are thought to be linked to emotional dysregulation observed in patients with MDD[45,47,48]. The visual regions involved in attention and perception further emphasize the multifaceted nature of MDD, where not only emotional processing but also sensory integration may influence treatment outcomes. The negative model weights observed for the frontoparietal and dorsal attention networks suggest that insufficient normalization of deviations within executive and attentional systems is linked to poorer treatment improvement in MDD. Prior morphometric studies have consistently reported reduced gray matter volume in frontal–parietal regions, with such alterations linked to cognitive dysfunctions and poorer clinical outcomes[7,49,50]. These findings indicate that disrupted FPN/DAN structure may serve as a neurobiological substrate for depression.

In contrast, Subtype 2 showed no significant prediction of HAMD change from GMV deviation shifts. Instead, in subtype 2, multivariate analysis revealed that changes in GMV deviations after treatment were systematically associated with baseline symptom severity, such that specific deviation-change patterns were linked to higher baseline HAMD-17 scores. This finding is compatible with previous studies. For instance, Mo et al. reported that specific GMV alterations (e.g. in supplementary motor area and occipital cortex) correlated with symptom dimensions (somatic complaints, alexithymia) at baseline in adolescent depression[51]. These observations suggest that subtype 2 may correspond to a more chronic or trait-like MDD phenotype: structural deviations reflect enduring illness severity rather than plasticity associated with short-term treatment gains.

In summary, the divergent longitudinal GMV changes of Subtype 1 and 2 illuminate the complex pathophysiology of MDD. In Subtype 1, restoration of specific GMV deficits (hippocampus, prefrontal/limbic regions) appears to correlate with symptom remission, whereas in Subtype 2 structural deviations seem proportionally linked to baseline severity. These findings reinforce the growing view that MDD is a biologically heterogeneous syndrome[4][41]. Mapping individual GMV deviation profiles, as demonstrated in our study, may provide a promising basis for predicting treatment responsiveness in specific patient subgroups and could ultimately contribute to more personalized intervention strategies.

### Limitations and Future Directions

Several limitations of this study should be acknowledged. First, we adopted a hard clustering algorithm (k-means), which assumes that each patient belongs to only one subtype. However, in reality, individuals may be influenced by multiple pathological processes simultaneously. Future work could employ soft clustering or latent variable models to allow for overlapping subtype features. Beyond GMV, future work could include resting-state fMRI, diffusion tensor imaging, and genetic or epigenetic information to construct multimodal normative models.

In summary, applying normative modeling to MDD subtyping holds great potential for revealing individual-level brain structural differences and advancing precision psychiatry, though further methodological refinement and clinical validation are essential.

## Declaration of interests

The authors declare no competing interests.

## Notes

### Competing Interest Statement

The authors have declared no competing interest.

